# IFNγ-associated immune–metabolic remodeling drives serotonin–kynurenine imbalance with cortical vulnerability in lupus-prone mice

**DOI:** 10.1101/2025.11.19.689278

**Authors:** Karim Matmat, Rosa-Maria Guéant-Rodriguez, Emmanuel Darcq, Okan Baspinar, Jean-Marc Alberto, Jean-Louis Guéant, Ayikoé-Guy Mensah-Nyagan, Hélène Jeltsch-David

## Abstract

**Introduction:** Neuropsychiatric systemic lupus erythematosus (NPSLE) is a major clinical challenge, characterized by heterogeneous manifestations and the absence of reliable biomarkers. The mechanisms linking systemic autoimmunity to neuronal injury and neuropsychiatric symptoms remain poorly understood.

**Methods:** Using the lupus-prone MRL/Lpr mouse model, we integrated systemic cytokine profiling, plasma neurofilament light chain (NfL), region-specific CNS cytokine mapping, cortical metabolomics, and behavioral analyses to dissect immune–metabolic–neuronal interactions.

**Results:** Inflammation was dominated by a Th1 cytokine program, with interferon-gamma emerging as the central driver. Composite cytokine scores correlated strongly with plasma NfL, establishing an immune–neuronal injury axis. Region-resolved analyses revealed distinct CNS cytokine signatures, including selective hippocampal loss of interleukin-10 and IFNγ-dominated responses in the frontal cortex. Cortical metabolomics demonstrated diversion of tryptophan metabolism away from serotonin toward the kynurenine pathway, with increased quinolinic acid/kynurenic acid (QA/KA) ratio and upregulation of indoleamine 2,3-dioxygenase-1 (*Ido1*) and kynurenine 3-monooxygenase (*Kmo*). NfL levels were negatively associated with serotonin and positively with 3-hydroxykynurenine and QA/KA, linking axonal damage to an excitotoxic metabolic environment. Importantly, cortical serotonin levels correlated with exploratory behavior, linking serotonergic depletion to anxiety-like phenotypes.

**Discussion:** Together, these results delineate a cascade in which systemic IFNγ is associated with cortical metabolic reprogramming and neuronal vulnerability, bridging peripheral immune activation with serotonergic depletion, melatonin loss, axonal injury, and behavioral dysfunction. Translationally, combined blood or CSF monitoring of IFNγ, NfL, and kynurenine metabolites could represent a candidate biomarker framework for NPSLE. However, validation in independent patient cohorts will be essential, and therapeutic modulation of IDO1/KMO or serotonergic pathways remains an avenue for future investigation.

## Introduction

Systemic lupus erythematosus (SLE) is a chronic autoimmune disease characterized by systemic inflammation, loss of immune tolerance, and involvement of multiple organ systems (1–3). Beyond its classic peripheral manifestations, SLE frequently presents with neurological and psychiatric symptoms, collectively termed “neuropsychiatric SLE” (NPSLE). These manifestations occur in a substantial proportion of patients, affecting ∼40% in prospective cohorts and up to 75% over the disease course (4,5). Clinical features include cognitive dysfunction, mood and anxiety disorders, seizures, cerebrovascular events, and psychosis (5). This clinical heterogeneity of NPSLE, combined with the lack of reliable biomarkers, poses significant challenges for diagnosis and patient stratification (6). Cognitive dysfunction, in particular, is among the most common manifestations but often remains underrecognized in routine clinical practice (7), highlighting the urgent need for translational, biomarker-based approaches.

Progress in understanding NPSLE has been hampered by the limited availability of human brain tissue and the rarity of longitudinal patient cohorts with well-characterized neuropsychiatric involvement (8). In this context, animal models play a critical role in elucidating disease mechanisms and identifying candidate biomarkers and therapeutic targets. Among the available models, the MRL/MpJ-Faslpr (MRL/Lpr) mouse is widely recognized as a reference model for lupus research (9–11). This strain carries a mutation in the Fas receptor gene (lpr), which impairs lymphocyte apoptosis and results in massive lymphoid hyperplasia, accumulation of double-negative T cells, and production of typical lupus autoantibodies, including antinuclear (ANA), anti–double-stranded DNA (anti-dsDNA), anti-Smith (anti-Sm), anti-Ro/SSA, and anti-La/SSB antibodies (12). As a consequence, MRL/Lpr mice spontaneously develop the hallmark features of systemic autoimmunity, such as immune complex deposition, vasculitis, and multi-organ inflammation. Importantly, they also exhibit well-documented neurobehavioral alterations, such as anxiety-like behavior, cognitive deficits, and depression-like phenotypes, accompanied by central nervous system (CNS) inflammation and metabolic changes (9,11,13,14). These features closely mirror key neuropsychiatric symptoms observed in SLE patients, providing a functional bridge between immune–metabolic dysregulation and CNS dysfunction. The combination of systemic autoimmunity, neuropathology, and behavioral outcomes makes the MRL/lpr strain a robust and translationally relevant model for investigating the pathogenesis of NPSLE.

Neuroinflammation is a central driver of CNS dysfunction, with peripheral cytokines known to disrupt blood–brain barrier integrity, activate microglia, and impair synaptic plasticity (15). In NPSLE, elevated levels of pro-inflammatory cytokines have been detected in both serum and cerebrospinal fluid (CSF); however, the precise relationship between these inflammatory mediators, neuronal injury, and cognitive outcomes remains incompletely understood (16,17). To date, most mechanistic studies in lupus-prone mice have relied on whole-brain homogenates, potentially masking region-specific alterations that are critical for understanding disease pathophysiology (11,18–20). Exploring regional vulnerability within the brain thus represents a crucial next step, as it may shed light on the diverse and heterogeneous neuropsychiatric manifestations observed in NPSLE patients (13,21). Equally important is the pressing need to identify peripheral biomarkers capable of reflecting such central alterations in a clinically accessible manner.

One potential mechanistic link between peripheral immune activation and brain dysfunction is tryptophan (TRP) metabolism. TRP is the precursor of serotonin, a neurotransmitter crucial for regulating mood, cognition, and executive processes via cortical and limbic circuits (22–24). Under inflammatory conditions, interferon-gamma (IFNγ) induces the enzyme indoleamine 2,3-dioxygenase-1 (IDO1), which diverts TRP catabolism away from serotonin biosynthesis toward the kynurenine pathway (KP). This shift results in the production of metabolites with opposing neuroactive properties: kynurenic acid (KA) exerts neuroprotective actions, whereas 3-hydroxykynurenine (3-HK) and quinolinic acid (QA) promote oxidative stress and excitotoxicity (25,26). Clinical studies have reported altered TRP metabolism in SLE patients, and similar alterations have been observed in the brains of MRL/Lpr mice (27). However, the mechanisms by which peripheral immune activation influences cortical vulnerability through the KP, and how these metabolic changes contribute to neuropsychiatric symptoms, remain poorly understood.

Beyond metabolic dysregulation, objective markers of neuronal injury are critical for assessing the impact of inflammation on the brain. One such marker, neurofilament light chain (NfL), has gained attention as a highly sensitive and reliable biomarker of axonal damage, detectable in blood with excellent analytical sensitivity (28). Elevated NfL levels have been reported in SLE patients, particularly in those with neuropsychiatric involvement, and show strong correlations with CSF concentrations (29–31). However, despite its validation in multiple sclerosis (MS) and Alzheimer’s disease (AD), NfL is not yet established in routine practice for lupus, and its capacity to detect cognitive or psychiatric involvement in NPSLE has yet to be fully characterized (32,33). Incorporating NfL measurements into preclinical lupus models offers a promising avenue to bridge this gap, enabling systematic integration with upstream inflammatory and metabolic markers, such as cytokines and tryptophan pathway metabolites, to generate a more comprehensive and translationally relevant profile of CNS involvement.

In this study, we implemented an integrated framework to examine immune activation, metabolic dysregulation, neuronal injury, and behavioral readouts in the MRL/Lpr lupus model. Peripheral inflammation was profiled using high-sensitivity multiplex immunoassays, with a focus on IFNγ, and combined with plasma NfL measurements. Notably, in this context, NfL elevation was not interpreted solely as an indicator of neuronal injury, but also as a downstream readout of inflammation-induced metabolic remodeling, specifically, shifts in the serotonin–kynurenine (KYN) balance. Importantly, by linking these molecular signatures to anxiety-like behavior and memory impairments, our framework offers a multidimensional perspective that connects systemic immune activation to cortical metabolic remodeling, neuronal damage, and functional impairment. We therefore propose that integrating inflammatory, metabolic, and neuronal injury markers can yield a clinically accessible and mechanistically informed strategy to detect brain alterations in NPSLE, with promising implications for future patient stratification and targeted therapeutic strategies.

## Materials and methods

### Animals, experimental design, and tissue collection

#### Animals, longitudinal monitoring, and sacrifice

Female MRL/MpJ-Faslpr (MRL/Lpr; stock #000485) and congenic MRL/MpJ controls (MRL^+/+^; stock #000486) mice were obtained from The Jackson Laboratory (Bar Harbor, ME, USA). Female mice were selected based on the higher prevalence of NPSLE in women and the consistent use of females in prior MRL/Lpr studies. Nevertheless, the absence of male cohorts limits generalization and precludes evaluation of potential sex-specific differences, future studies will be required to address this gap. Animals were housed in individually ventilated cages (four mice per cage) under standard laboratory conditions (22 ± 1 °C; 12-h light/dark cycle), with *ad libitum* access to food and water. Body weight was monitored weekly from 4 to 17 weeks of age, covering the full experimental timeline from arrival to sacrifice. As a non-invasive readout of renal disease progression, proteinuria was assessed at regular intervals using Albustix^®^ reagent strips (Siemens). At 17 weeks of age, corresponding to the peak of systemic and neuropsychiatric manifestations in MRL/Lpr mice, animals were anesthetized with isoflurane (4% for induction in an induction chamber and 2% for maintenance in oxygen, delivered via a precision vaporizer) and euthanized by decapitation while under deep anesthesia. Immediately after euthanasia, the kidneys, brain, spinal cord, and spleen were rapidly dissected and weighed.

All procedures complied with the European Directive 2010/63/EU and the French governmental decree 2013-118. Experimental protocols were approved by the local Animal Ethics Committee (APAFIS#35144) and conducted under the supervision of authorized investigators in a certified animal facility (Faculty of Medicine, University of Strasbourg).

#### Brain and spinal cord processing

Brains were rapidly removed on ice and bisected along the mid-sagittal plane. One hemisphere was preserved intact for whole-brain analyses, whereas the contralateral hemisphere was microdissected to isolate the frontal cortex, hippocampus, and cerebellum. Samples were snap-frozen in liquid nitrogen and stored at −80 °C. The spinal cord was isolated by hydraulic extrusion: A 10-mL syringe filled with ice-cold PBS (pH 7.4) was inserted into the sacral/lumbar canal, and gentle pressure was applied to flush the spinal cord out through the cranial opening. The entire cord (cervical, thoracic, lumbar) was collected, snap-frozen, and stored at −80 °C.

#### Blood collection and plasma preparation

During decapitation, blood was collected into EDTA-coated tubes kept on ice. Plasma was separated by centrifugation at 2,000 × g for 10 min at 4 °C, aliquoted, and stored at −80 °C until further use.

### Behavioral Testing

Behavioral evaluation was conducted sequentially in the same cohort of animals: the Y-Maze spontaneous alternation task at 14 weeks of age, the Novel Object Recognition (NOR) test at 15 weeks of age, and the Open Field (OF) test at 16 weeks of age (34). Tests were spaced at one-week intervals to allow sufficient recovery and to minimize carryover effects. All tests were conducted under stable ambient illumination (50 lux).

#### Y-Maze spontaneous alternation

Spontaneous alternation behavior (SAB), a measure of spatial working memory, was assessed using a Y-maze made of opaque grey Perspex with three identical arms (35 × 7 × 15 cm) positioned at 120° angle. Each mouse was placed in the central zone and allowed to freely explore all three arms for 6 min. SAB reflects the natural tendency of rodents to alternate between arms during exploration. Arm entries were automatically recorded, and alternation sequences were defined as three consecutive entries into each of the three arms without repetition. The SAB score was calculated as:

% Alternation = Number of Alternations / (Total Arm Entries -2) x100.

The total number of arm entries was also measured as an index of locomotor activity (data not shown).

#### Novel Object Recognition (NOR)

NOR was assessed in transparent Plexiglas arenas (40 × 40 × 40 cm). The task was performed over three consecutive days. On day 1 (habituation phase), mice were placed in the empty arena to acclimate to the environment. On day 2 (spatial habituation phase), three identical objects (green plastic capsules of identical size, shape, and color) were positioned equidistantly within the same arena, and mice were allowed to explore freely for 10 min, establishing familiarity with the objects. On day 3 (recognition test), one of the familiar objects was replaced with a novel object of similar size but different shape and color. Each session lasted 10 min. Exploration was defined as direct head contact with the object. The arena and all objects were cleaned with 70% ethanol between trials to eliminate olfactory cues. Exploration time for each object was quantified, and the recognition index (RI) was calculated as:

RI (%) = (T_novel_) / (T_novel_ + T_Familiar1_ + T_Familiar2_) x 100, where T_novel_ is the exploration time of the novel object and T_Familiar1,2_ are the exploration times of the two remaining familiar objects. A higher RI reflects improved memory recognition performance. Behavioral tracking and analysis were performed using AnyMaze software v7.4

#### Open Field (OF)

The OF test was conducted to assess general locomotor activity and anxiety-like behavior. Mice were placed individually into the center of a square open field arena (40 × 40 × 40 cm) and allowed to explore freely for 15 minutes. The arena was evenly illuminated (50 lux) and cleaned with 70% ethanol between trials to eliminate olfactory cues. The behavior was assessed using the automated SuperFlex Open Field system (Omnitech Electronics, Columbus, USA), which consists of six independent arenas equipped with infrared beam tracking. For analysis, each arena was virtually divided into three zones: (i) the central zone (anxiety-relevant open area), (ii) the peripheral zone, and (iii) the corner zones, representing the areas closest to the walls. Infrared beam breaks were recorded and analyzed by the Fusion SuperFlex Edition software *v6.5* (Omnitech), providing automated tracking of locomotor activity and zone occupancy. Anxiety-related behavior was evaluated based on the time spent in the central zone, the number of entries into the central zone, the distance traveled in the central zone, and the percentage of central distance relative to the total distance traveled (34). To ensure that anxiety measures were not confounded by motor impairments, overall locomotor activity was assessed by the total distance traveled and the ambulatory time.

### Plasma Cytokines and Neurofilament Light Chain (NfL) Quantification by MSD U-PLEX

Plasma concentrations of interleukin-1 beta (IL-1β), interleukin-6 (IL-6), interleukin-10 (IL-10), granulocyte–macrophage colony-stimulating factor (GM-CSF), interleukin-17A (IL-17A), tumor necrosis factor alpha (TNFα), and IFNγ were quantified using a custom U-PLEX multiplex assay (Meso Scale Discovery, MSD, USA). NfL levels were measured separately using a U-PLEX singleplex assay (MSD, USA). All assays were performed on EDTA plasma samples according to the manufacturer’s instructions.

Briefly, plasma samples (MRL^+/+^, n = 13; MRL/Lpr, n = 12) were thawed on ice, centrifuged at 2,000 × g for 10 min at 4 °C to remove debris, and diluted in assay diluent as recommended. Standards and samples (50 µL/well, run in triplicate) were loaded onto pre-coated plates and incubated at room temperature with shaking. Plates were then washed and incubated with SULFO-TAG–labeled detection antibodies. Following a final wash, MSD Read Buffer was added, and electrochemiluminescence (ECL) signals were immediately acquired using a MESO QuickPlex SQ 120 instrument (MSD, USA).

Analyte concentrations were calculated from standard curves generated with MSD Discovery Workbench software v4.0 and expressed as pg/mL. Samples with values falling outside the reliable range of the standard curve were excluded from the analyses.

### RNA Isolation Quantitative and RT-PCR Analysis

Total RNA was extracted from frozen tissues, including whole brain, frontal cortex, hippocampus, cerebellum, and spinal cord, using the NucleoSpin® RNA/Protein kit (Macherey-Nagel, Germany), according to the manufacturer’s instructions. For each region and animal (MRL^+/+^, n = 4–5; MRL/Lpr, n = 4–5), 500 ng of total RNA were used for two-step reverse transcription quantitative PCR (RT-qPCR). Group sizes were necessarily limited by the multi-regional dissection design. While this reduces statistical power, the consistency of gene expression trends across animals, and their convergence with metabolomic and cytokine findings support the robustness of the conclusions.

Reverse transcription was performed using PrimeScript™ RT Master Mix (Takara, Japan), and qPCR was carried out with SYBR® Premix Ex Taq™ (Takara) as previously described (35). Specific primers (Supplementary Material S1) were synthesized by Eurogentec (Angers, France). Cycle threshold (Ct) values were determined for each reaction, and relative expression levels of target genes were normalized to the housekeeping genes *Hprt1* and *Polr2a* using the 2^−ΔΔCt^ method. The stability of housekeeping gene expression across tissue types was systematically assessed using the gene stability module of Bio-Rad CFX software v3.0, ensuring appropriate normalization.

### Targeted Metabolite Quantification by LC-MS/MS Analyses

Targeted metabolite analysis was performed on frontal cortex tissue using liquid chromatography–tandem mass spectrometry (LC-MS/MS). Samples (MRL^+/+^, n = 13; MRL/Lpr, n = 12) were homogenized in 1× phosphate-buffered saline (PBS) using microtubes containing ceramic beads. Homogenates were centrifuged at 12,000 × g for 20 min at 4 °C, and the resulting supernatants were collected for further processing. Supernatants were spiked with internal standards, including dithiothreitol (DTT), to control for variability in sample processing. Proteins were precipitated by the addition of cold methanol, followed by a 30-minute incubation on ice. Samples were then diluted fourfold with 0.1% formic acid solution. Metabolite quantification was performed on a Shimadzu LCMS-8045 electrospray ionization triple quadrupole system (Shimadzu, Kyoto, Japan). Data acquisition and processing were carried out using Insight software (v3.1, Shimadzu). To ensure data reliability, samples with concentrations falling near the instrumental detection limits were excluded from the analysis, and only values within the validated detection range were retained. Final metabolite concentrations were normalized to the total protein concentration of each sample, determined using the bicinchoninic acid (BCA) assay (Interchim, France).

### Statistical and Computational Analysis

#### Software Environment

All statistical analyses were performed in R v4.3.0 within the RStudio 2025.05.0 Build 496 environment. Core functions from the *stats* package were used for hypothesis testing and regression modeling.

#### Data Visualization

Data were visualized using ggplot2, ggrepel, and other relevant R packages. Specifically: Behavioral outcomes, metabolite concentrations, and metabolite ratios were visualized as dot plots showing individual values overlaid with group mean ± standard error of the mean (SEM). NfL levels and the composite inflammatory score were displayed as boxplots indicating distribution from minimum to maximum with the median. RT-qPCR data were represented as bar plots showing mean ± SEM. Correlation analyses were visualized as scatter plots with regression lines and 95% confidence intervals. Multivariate analyses included heatmaps (*ComplexHeatmap*), radar plots (*fmsb*), and principal component analysis (PCA) biplots generated with prcomp on centered and scaled data. PCA plots included both sample scores and variable loadings, and visualization was performed with *ggplot2* and *ggrepel*. Schematic illustrations and graphical summaries were created with BioRender.com.

#### Group Comparisons

Normality was assessed with the Shapiro–Wilk test and homogeneity of variance with the F test. Depending on distribution, between-group comparisons were made using two-tailed unpaired Student’s t-tests (parametric), Welch’s t-tests (for unequal variance), or Mann–Whitney U tests (non-parametric). A significance threshold of *P* < 0.05 was applied. For analyses involving multiple comparisons (*e.g.*, metabolite panels and correlations), p-values were adjusted using the Benjamini–Hochberg false discovery rate (FDR) procedure, and adjusted values (*q*) are reported.

#### Behavioral Data Analysis

For behavioral tests (Open Field, Y-Maze, Novel Object Recognition), comparisons between groups were made using the same statistical pipeline (*e.g.*, *Group comparisons section*). Group sizes ranged between 10 and 13 animals per genotype, as some mice were excluded due to technical issues (*e.g.*, recording failures) or non-responsiveness (*e.g.*, no exploration during NOR, total immobility during OF/Y-Maze). This ensured unbiased statistical analyses and interpretable results.

#### Correlation Analyses

Correlations between matched biological and behavioral parameters were assessed using Pearson’s or Spearman’s correlation coefficients, depending on data distribution. Robustness was assessed with a leave-one-out procedure, where Δρ quantified the maximum change in correlation upon sequential removal of each sample. Multiple testing was controlled with Benjamini–Hochberg FDR correction.

#### Receiver Operating Characteristic (ROC) analysis

ROC analyses were performed in R using the *pROC* package to evaluate the discriminative performance of the composite inflammatory score derived from the z-scored plasma levels of seven cytokines (IFNγ, TNF-α, IL-1β, IL-6, IL-10, IL-17A, GM-CSF). ROC curves were generated by plotting sensitivity against 1−specificity, and the area under the curve (AUC) was computed with 95% confidence intervals obtained by bootstrap resampling (2000 iterations). Optimal thresholds were determined using Youden’s index, and the corresponding sensitivity and specificity values are reported.

## Results

### Systemic and Behavioral Characterization of MRL/Lpr Lupus Model

To establish baseline systemic and behavioral features of the MRL/Lpr lupus model, we longitudinally monitored body weight and proteinuria from 4 to 17 weeks of age. Body weight increased steadily over time in both MRL/Lpr and MRL^+/+^ mice with no significant differences between groups (Fig. 1A). In contrast, proteinuria rose markedly in MRL/Lpr mice from 11 weeks onward, reaching high levels by 17 weeks (Fig. 1B). This renal dysfunction was further supported by a significant increase in kidney weight at sacrifice (*P* < 0.0001; Fig. 1C), consistent with functional renal impairment and indicative of lupus nephritis. Splenomegaly was also pronounced in MRL/Lpr mice relative to controls (*P* < 0.0001; Fig. 1D), reflecting systemic autoimmunity and chronic immune activation, hallmarks of the model. By contrast, brain weight was unchanged (Fig. 1E), suggesting preserved global brain mass, while spinal cord weight was significantly increased in MRL/Lpr mice (*P* = 0.0043; Fig. 1F). Although the underlying mechanism remains unclear, this enlargement may reflect inflammatory infiltration or gliotic changes, but histopathological confirmation is needed.

**Figure 1:**
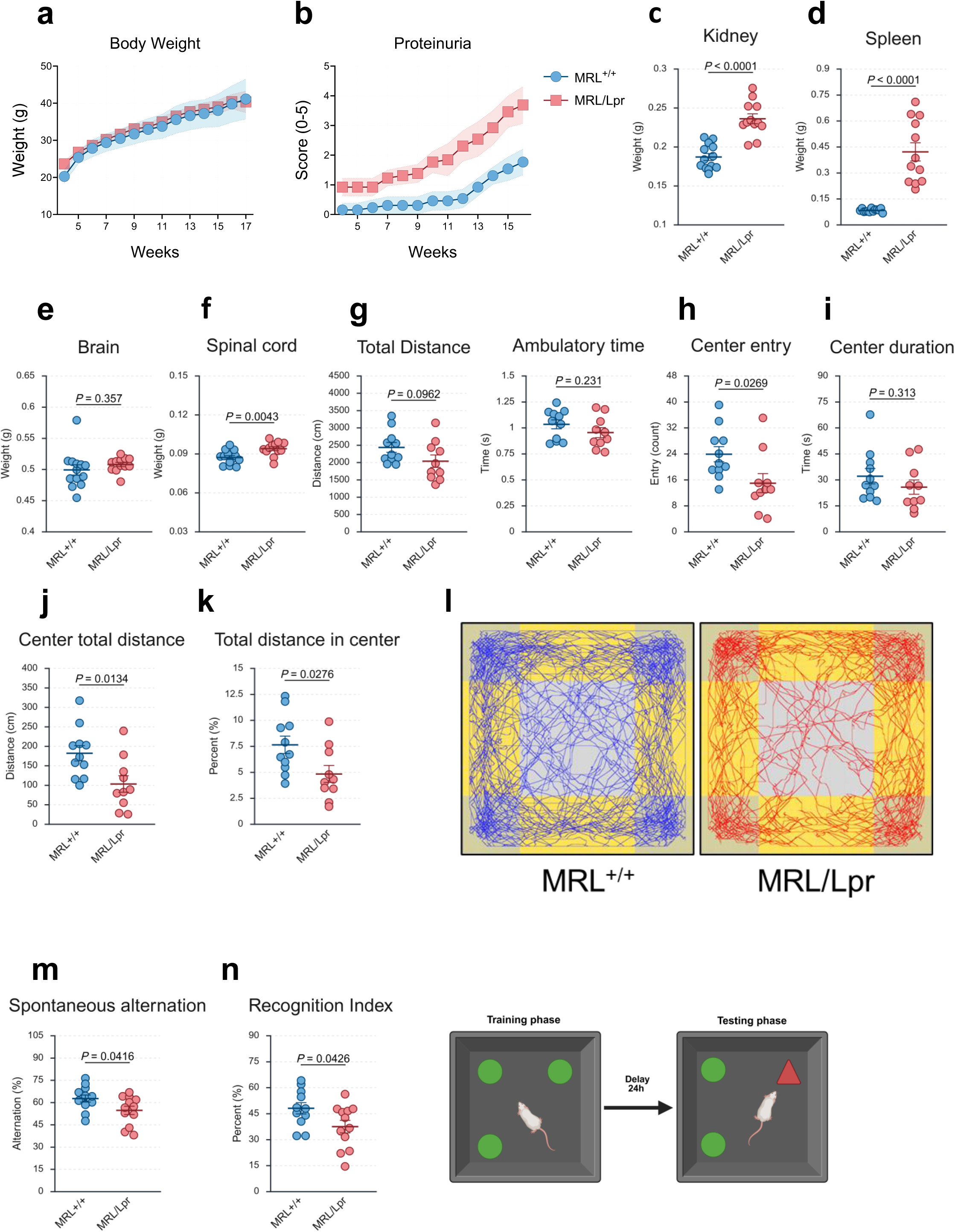
Systemic autoimmunity in MRL/Lpr mice associates with anxiety-like behavior and cognitive deficits. **(A)** Body weight progression from 4 to 17 weeks shows no significant differences between MRL^+/+^ and MRL/Lpr mice. Shaded areas represent 95% confidence intervals. **(B)** Proteinuria scores increase markedly in MRL/Lpr mice from 11 weeks onward, indicating renal dysfunction. Shaded areas represent 95% confidence intervals. **(C–F)** Organ weights at 17 weeks of age: MRL/Lpr mice exhibit nephromegaly (*P* < 0.0001; C), splenomegaly (*P* < 0.0001; D), preserved brain weight (E), and increased spinal cord weight (*P* = 0.0043; F), consistent with systemic inflammation and potential CNS involvement. **(G)** Locomotor activity in the OF test: total distance and ambulatory time are comparable between groups. **(H-K)** Anxiety-like behavior in the OF test: MRL/Lpr mice show fewer entries into the center (*P* = 0.0269; H), no significant difference in time spent in the center (*P* = 0.313; I), reduced distance traveled in the center (*P* = 0.0134; J), and a lower proportion of center exploration (*P* = 0.0276; K), **(L)** Representative activity tracks in the OF from three mice per group illustrate lack of consistent central exploration in MRL/Lpr mice. (**M-N**) Cognitive performance: MRL/Lpr mice display reduced spontaneous alternation in the Y-maze (*P* = 0.0416; M) and a lower recognition index in the NOR task (*P* = 0.0426; N), indicating deficits in spatial working memory and recognition memory. Locomotor parameters were comparable between groups across tests. Data are shown as mean ± SEM*. n* = 10–13 mice per group. Group comparisons were performed using appropriate parametric or non-parametric tests depending on data distribution (see Methods for details). *P* < 0.05 considered statistically significant.

Total distance and ambulatory time were comparable between groups (Fig. 1G), excluding major motor impairments and ensuring that subsequent measures were not confounded by differences in general activity. However, MRL/Lpr mice exhibited clear anxiety-like behavior, with fewer entries into the center (*P* = 0.0269; Fig. 1H), while the total time spent in the center did not differ significantly (Fig. 1I). The distance traveled in the center was also reduced (*P* = 0.0134; Fig. 1J), with a lower proportion of central exploration relative to total distance (*P* = 0.0276; Fig. 1K). The preserved total time spent in the center likely reflects variability from occasional prolonged visits. Representative trajectories further highlight the lack of consistent exploration of the central zone in MRL/Lpr mice compared with controls (Fig. 1L).

Cognitive performance was evaluated using the Y-maze and NOR tasks. MRL/lpr mice exhibited reduced spontaneous alternation (*P* = 0.0416; Fig. 1M), indicating impaired spatial working memory, and a significantly lower recognition index in the NOR task (*P* = 0.0426; Fig. 1N). During the training phase, both groups explored identical objects equally (data not shown), confirming comparable baseline exploratory drive. At test, control MRL^+/+^ mice displayed a clear preference for the novel object, whereas MRL/lpr mice did not, consistent with a deficit in recognition memory. Importantly, locomotor activity during both tasks, including total distance, ambulatory time, and Y-maze entries, was comparable across groups, ruling out motor confounds.

Together, these results demonstrate that MRL/lpr mice display the hallmark systemic features of lupus, including marked splenomegaly and lymphadenopathy, alongside central alterations such as spinal cord hypertrophy, memory impairments, and increased anxiety-like behavior.

### Peripheral Cytokine Upregulation Defines a Systemic Inflammatory Signature Associated with Neuronal Injury

Building on the systemic and behavioral characterization, we investigated whether MRL/Lpr mice display a distinct peripheral inflammatory profile and whether this relates to neuroaxonal injury.

Plasma cytokine levels were quantified using a multiplex Meso Scale Discovery (MSD) U-PLEX assay. A global overview of the cytokine profile, visualized as a radar plot (Fig. 2A), revealed a robust pro-inflammatory signature in MRL/Lpr mice compared with MRL^+/+^ controls, characterized by marked upregulation of IFNγ, TNFα, IL-6, and IL-1β. IL-10 was also elevated, consistent with its complex role in lupus. In contrast, IL-17A and GM-CSF remained unchanged, despite their well-established roles in T cell–driven inflammation. To assess the coordinated structure of the cytokine data, we performed principal component analysis (PCA). The first principal component (PC1) accounted for ∼49% of the total variance and was driven primarily by IFNγ, IL-10, and TNFα (Fig. 2B). PCA confirmed a clear separation between MRL/Lpr and control animals, indicating a genotype-specific systemic cytokine program.

**Figure 2:**
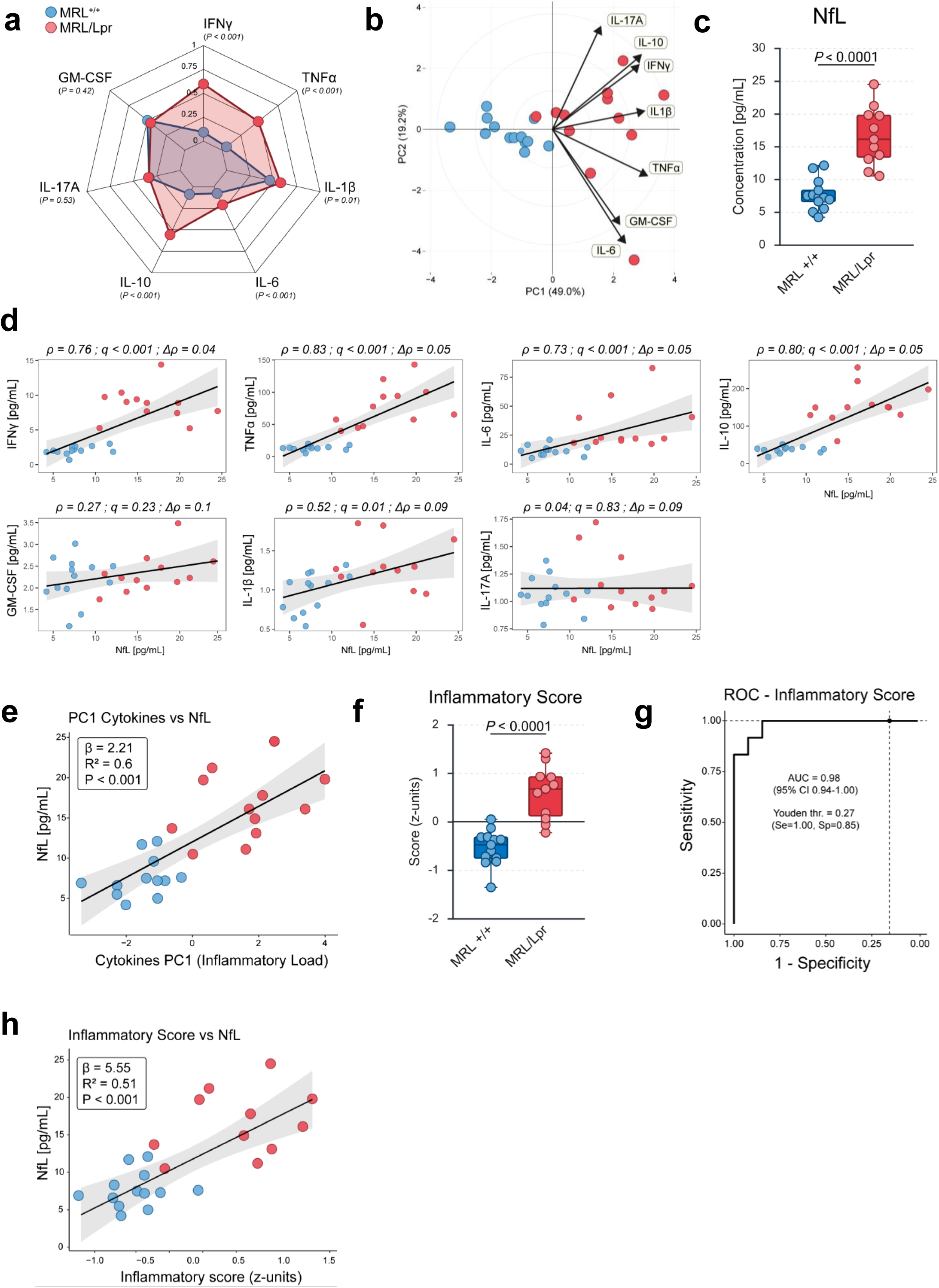
A Th1-dominated cytokine program drives plasma NfL elevation, linking inflammation to neuronal injury. **(A)** Radar plot of plasma cytokines in MRL^+/+^ and MRL/Lpr mice. MRL/Lpr mice exhibit a robust pro-inflammatory signature, characterized by elevated levels of IFNγ, TNFα, IL-6, IL-1β, and IL-10. IL-17A and GM-CSF levels remained unchanged. Data are expressed as normalized z-scores (0–1) computed from raw concentrations (pg/mL). *P* values for group comparisons are indicated on the plot. **(B)** Principal component analysis (PCA) of plasma cytokine concentrations, showing separation of MRL^+/+^ and MRL/Lpr mice along PC1 (49.0% variance explained) and PC2 (19.2%). Loading vectors indicate the contribution of individual cytokines to each component. **(C)** Plasma neurofilament light chain (NfL) concentrations (pg/mL) in MRL^+/+^ and MRL/Lpr mice. **(D)** Correlation analysis showing positive associations between plasma NfL levels and individual cytokines. Robustness of correlations was confirmed via leave-one-out analysis (Δρ < 0.2). Scatter plots display regression lines with shaded 95% confidence intervals. For each panel, Spearman’s or Pearson’s correlation was applied depending on data distribution, with false discovery rate (FDR) correction for multiple testing. ρ and Δρ values are indicated on the plots. **(E)** PC1 scores from the cytokine PCA strongly correlated with plasma NfL levels (R² ≥ 0.5, *P* < 0.001). **(F)** Inflammatory score computed as the mean of z-scores across all cytokines, compared between MRL^+/+^ and MRL/Lpr mice, providing an index of systemic inflammatory burden. **(G)** Receiver operating characteristic (ROC) curve of the inflammatory score for discriminating MRL^+/+^ and MRL/Lpr mice, with area under the curve (AUC), 95% confidence interval, and optimal Youden threshold (Se, Sp) indicated. **(H)** Correlation between plasma NfL and the composite inflammatory score (mean of cytokine z-scores). The composite inflammatory score showed a strong positive correlation with plasma NfL (R² ≥ 0.5, *P* < 0.001). Linear regression analysis is shown with slope (β), R², and *P* value. Each point represents an individual mouse, reflecting the sample size per group. Boxplots display median and range. Statistical analyses used parametric or non-parametric tests as appropriate (see Methods for details). *P* < 0.05 considered statistically significant.

To determine whether this systemic inflammatory profile was linked to neuronal injury, we quantified plasma NfL, a sensitive biomarker of axonal damage. NfL levels were markedly elevated in MRL/Lpr mice compared with controls (Fig. 2C). Correlation analyses revealed strong positive associations between NfL and IFNγ, IL-6, TNFα, IL-10, and IL-1β (all ρ ≥ 0.50, *q* < 0.05; Fig. 2D). Robustness of these relationships was confirmed by leave-one-out analysis (Δρ < 0.2), excluding the possibility that single outliers accounted for the observed effects. By contrast, IL-17A and GM-CSF showed no significant associations with NfL.

Because cytokines act synergistically, we next derived integrative indices of inflammatory burden. Building on the PCA results, we found that PC1 values correlated strongly with plasma NfL concentrations (R² ≥ 0.5, *P* < 0.001; Fig. 2E), highlighting that neuroaxonal injury is significantly explained by the coordinated cytokine program. As a complementary approach, we constructed a composite inflammatory score by standardizing each cytokine concentration (z-scores) and averaging them into a single value (Fig. 2F). This unweighted composite provides a simple, reproducible index of inflammatory load. Alternative approaches, such as PCA-weighted scoring, yielded highly similar results (data not shown), underscoring the robustness of this index. We then assessed its discriminative performance using ROC analysis (Fig. 2G). This approach evaluates whether the score can reliably classify mice according to genotype, quantifying sensitivity and specificity across all thresholds. The composite score achieved very high accuracy (AUC = 0.98, 95% CI: 0.94–1.00; Youden threshold = 0.27). Finally, the score correlated strongly with plasma NfL (R² ≥ 0.5, *P* < 0.001; Fig. 2H), supporting the view that aggregated cytokine activity provides a biologically meaningful representation of inflammatory burden associated with neuronal injury.

Collectively, these findings demonstrate that MRL/Lpr mice show a coordinated pro-inflammatory program dominated by IFNγ, TNFα, IL-6, and IL-10, which is tightly linked to biomarkers of neuronal injury. Integrative inflammatory indices, validated through both correlation with NfL and high ROC classification performance, provide a quantitative framework to capture the pathogenic bridge between systemic inflammation and neuronal damage in this lupus model.

### Central Inflammatory Responses Reveal Region-Specific Cytokine Patterns

Given the systemic inflammatory program observed in MRL/Lpr plasma, we next examined whether peripheral inflammation was mirrored within the CNS and whether specific brain regions displayed distinct inflammatory signatures. To address this, we quantified mRNA expression of key cytokines (*e.g.*; IL-1β, IL-6, IFNγ, and IL-10) in whole-brain homogenates and in dissected regions (*e.g.*; frontal cortex, hippocampus, cerebellum, and spinal cord). This design allowed us to capture not only global CNS dynamics but also potential region-specific heterogeneity.

A global overview visualized as a heatmap revealed a heterogeneous distribution of cytokine expression across regions in MRL/Lpr mice (Fig. 3A), pointing to marked spatial variability in central inflammatory responses. Consistent with plasma findings, whole-brain lysates confirmed that MRL/Lpr mice exhibited a global increase in pro-inflammatory mediators compared to controls (Fig. 3B). At the regional level, however, cytokine expression varied substantially. Log₂ fold-change analysis revealed a mosaic-like landscape: IL-1β induction was strong in the hippocampus and cerebellum, IL-6 was primarily elevated in the spinal cord and cerebellum, whereas IFN-γ emerged as the predominant signal in the frontal cortex and spinal cord (Fig. 3C). Interestingly, IL-10 displayed opposite trends depending on the region, being reduced in the hippocampus despite its increase elsewhere. Such divergence in IL-10 expression points to region-specific immune regulation, which may underlie differential vulnerability to inflammation-associated damage.. Finally, PCA of regional cytokine profiles (Fig. 3D) captured this heterogeneity at a systems level, showing clear segregation of brain regions into distinct clusters, each defined by different cytokine drivers. Indeed, while all four cytokines mRNA level contributed to the total variance, their influence was clearly region-dependent, supporting the existence of anatomically compartmentalized inflammatory programs.

**Figure 3:**
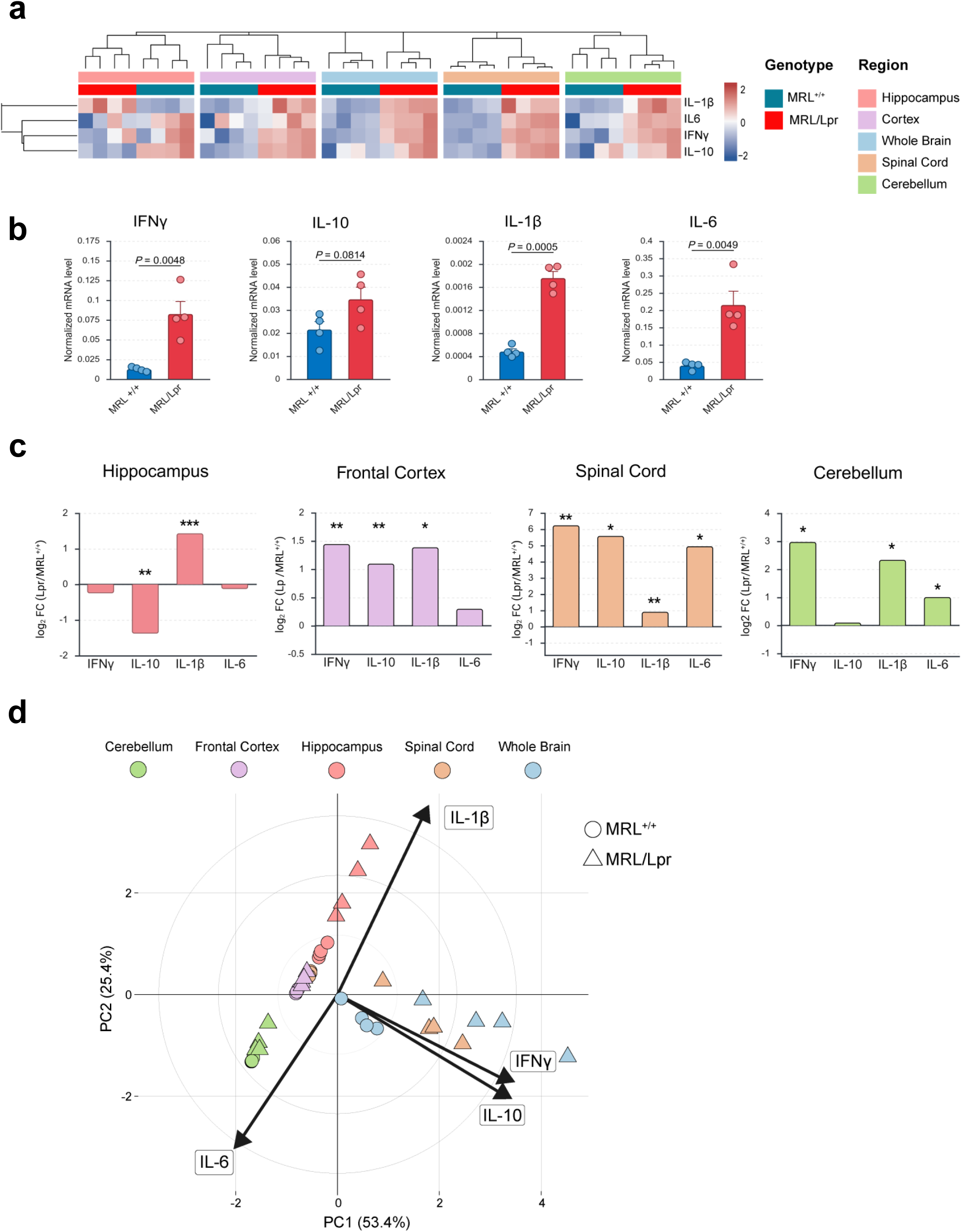
Central inflammation follows a mosaic pattern with frontal cortex IFNγ dominance and hippocampal IL-10 loss. **(A)** Hierarchical clustering heatmap of normalized mRNA expression levels (qPCR) of IL-1β, IL-6, IFNγ, and IL-10 across brain regions (frontal cortex, hippocampus, cerebellum, spinal cord) and whole brain homogenates in MRL^+/+^ and MRL/Lpr mice. MRL/Lpr mice exhibit a heterogeneous distribution of cytokine expression, indicating region-specific central inflammatory responses. **(B)** Normalized whole brain homogenate mRNA expression levels of IL-1β, IL-6, IFNγ, and IL-10. MRL/Lpr mice show a significant global increase in pro-inflammatory cytokines compared to MRL^+/+^ controls. **(C)** Regional expression patterns represented as Log2 fold-change (MRL/Lpr relative to MRL^+/+^) for IFNγ, IL-10, IL-1β, and IL-6, highlighting region-specific differences across CNS compartments. *Statistical significance is indicated by asterisks (\**P* < 0.05; \*\**P* < 0.01; \*\*\**P* < 0.001). **(D)** Principal component analysis (PCA) of regional cytokine expression profiles, showing both separation between MRL^+/+^ (circles) and MRL/Lpr (triangles) and, more prominently, clustering according to brain region (cerebellum, frontal cortex, hippocampus, spinal cord, and whole brain). Arrows indicate the loading vectors for IL-1β, IL-6, IFNγ, and IL-10. Each point in quantitative plots represents an individual mouse, reflecting the number of biological replicates per group. Bars indicate group mean ± SEM. Statistical analyses were performed using appropriate parametric or non-parametric tests depending on data distribution (see Methods for details). *P* < 0.05 considered statistically significant.

Together, these findings demonstrate that central inflammatory responses in MRL/Lpr mice mirror the systemic inflammatory signature but follow a mosaic-like, region-specific organization rather than a uniform activation pattern. Notably, the spinal cord displayed a strong pro-inflammatory profile consistent with its increased mass, while the hippocampus combined IL-1β induction with loss of IL-10, highlighting a potential mechanism of selective vulnerability. In the frontal cortex and spinal cord, IFNγ emerged as a dominant driver, reflecting the peripheral cytokine profile and raising the possibility of downstream metabolic reprogramming through induction of the KP. Because this pathway directly competes with serotonin synthesis for tryptophan availability, cortical IFNγ-driven inflammation may represent a critical link between peripheral immune activation and central serotonin/KYN imbalance, thereby contributing to the anxiety- and memory-related alterations observed in MRL/Lpr mice.

### Frontal Cortex Metabolic Profiling Reveals a Shift from Serotonin to Kynurenine Pathway Activity

Given the cortical dominance of IFNγ identified in the inflammatory cytokine profiling, and its well-established role as an inducer of IDO1, we next examined whether TRP metabolism was reprogrammed in the frontal cortex. This region was also chosen because of its central role in mood regulation, cognition, affective behavior, and its importance as a hub of monoaminergic metabolism. TRP can be processed along two major routes: conversion into serotonin via the serotonin pathway, which supports neurotransmission and neuroplasticity, or degradation through the KP, which yields a spectrum of metabolites with either neuroprotective (*e.g.*, KA) or neurotoxic (*e.g.*, QA; 3-HK) properties (Fig. 4A).

**Figure 4:**
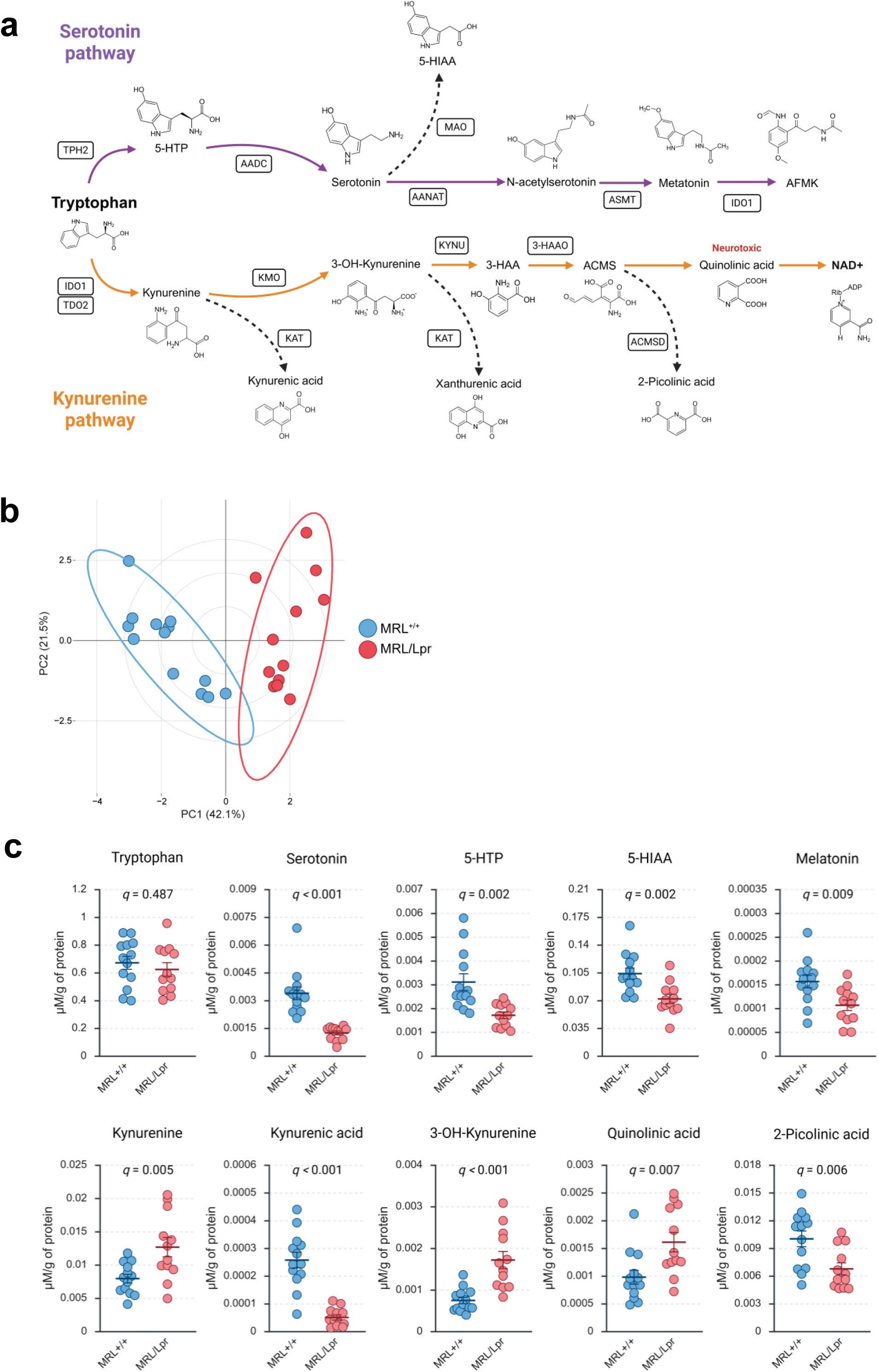
Frontal cortex tryptophan metabolism is shifted from the serotonin pathway toward kynurenine excitotoxicity. **(A)** Schematic overview of the serotonin (purple) and kynurenine (orange) pathways, indicating the enzymatic steps and their associated metabolites. Created with BioRender **(B)** Principal component analysis (PCA) of cortical tryptophan metabolites, showing clear separation between MRL^+/+^ and MRL/Lpr mice along PC1 (42.1% of variance) and PC2 (21.5%). **(C)** Quantification of individual tryptophan-derived metabolites in the frontal cortex, normalized to protein concentration (µM/µg protein). Each point represents an individual mouse, with bars indicating group mean ± SEM. Statistical analyses were performed using appropriate parametric or non-parametric tests depending on data distribution (see Methods for details). Multiple comparisons were corrected using the Benjamini–Hochberg false discovery rate (FDR) method, and a *q* value < 0.05 was considered statistically significant.

Targeted metabolomic profiling of the frontal cortex revealed a profound shift in TRP fate in MRL/Lpr mice compared with controls. PCA of metabolite profiles (Fig. 4B) demonstrated a clear separation between groups, indicating that the changes reflect a coordinated TRP metabolism reprogramming rather than isolated changes in individual compounds. Quantitative analyses (Fig. 4C) showed that, while TRP availability itself was unchanged, its metabolic rate was strongly diverted from serotonin biosynthesis toward the KP. Within the serotonin pathway, MRL/Lpr mice exhibited significant reductions in 5-hydroxytryptophan (5-HTP), the immediate precursor of serotonin, alongside decreased serotonin and its primary catabolite 5-hydroxyindoleacetic acid (5-HIAA). Levels of melatonin, a circadian regulator with antioxidant and neuroprotective functions, were also decreased, further indicating broad suppression of serotonergic metabolism. In contrast, KP metabolites displayed consistent upregulation. KYN, the central branching metabolite, accumulated significantly, indicating enhanced diversion of TRP into this pathway. Its downstream metabolism was oriented predominantly toward neurotoxic outputs, with elevated levels of 3-HK, a pro-oxidant metabolite that amplifies oxidative stress, and QA, an NMDA receptor agonist with excitotoxic and pro-inflammatory properties. By contrast, KA, a neuroprotective metabolite, was significantly reduced, further exacerbating the imbalance between neurotoxic and neuroprotective KP products. In addition, 2-picolinic acid (PA), a side branch product with reported neuroprotective functions, was also diminished.

Altogether, these data indicate a redirection of cortical tryptophan metabolism away from the serotonin pathway and toward a KP-biased, neurotoxic profile. This metabolic reprogramming is consistent with IFNγ-driven IDO1 activation and provides a mechanistic link between cortical inflammation, serotonin/KYN imbalance, and the anxiety- and memory-related alterations observed in MRL/Lpr mice. To further explore the regulatory basis of this metabolic remodeling, we next assessed the transcriptional regulation of key enzymes controlling TRP catabolism.

### Inflammation Diverts Tryptophan Metabolism Toward Excitotoxicity and Correlates with Neuroaxonal Injury

To obtain a functional readout of pathway activity, we first analyzed biologically relevant metabolite ratios in the frontal cortex (Fig. 5A–B). MRL/Lpr mice exhibited a marked increase in the KYN/TRP ratio, consistent with enhanced IDO1/TDO2 activity. The 3-HK/KYN ratio was also elevated, suggesting increased flux through kynurenine 3-monooxygenase (KMO) toward neurotoxic arm of the KP. In contrast, the 5-HTP/TRP ratio was reduced, reflecting impaired serotonin biosynthesis, while the 5-HIAA/serotonin ratio was elevated, consistent with accelerated serotonin catabolism. Ratios indexing the balance between neurotoxic and neuroprotective outputs were also altered: the QA/KA ratio was significantly increased, pointing to a shift toward excitotoxicity, whereas PA/QA was reduced, indicating a lack of compensatory engagement of protective branches. Together, these ratios consolidate the notion of a metabolic reprogramming that diverts TRP flux away from serotonin and toward the neurotoxic arm of the KP.

**Figure 5:**
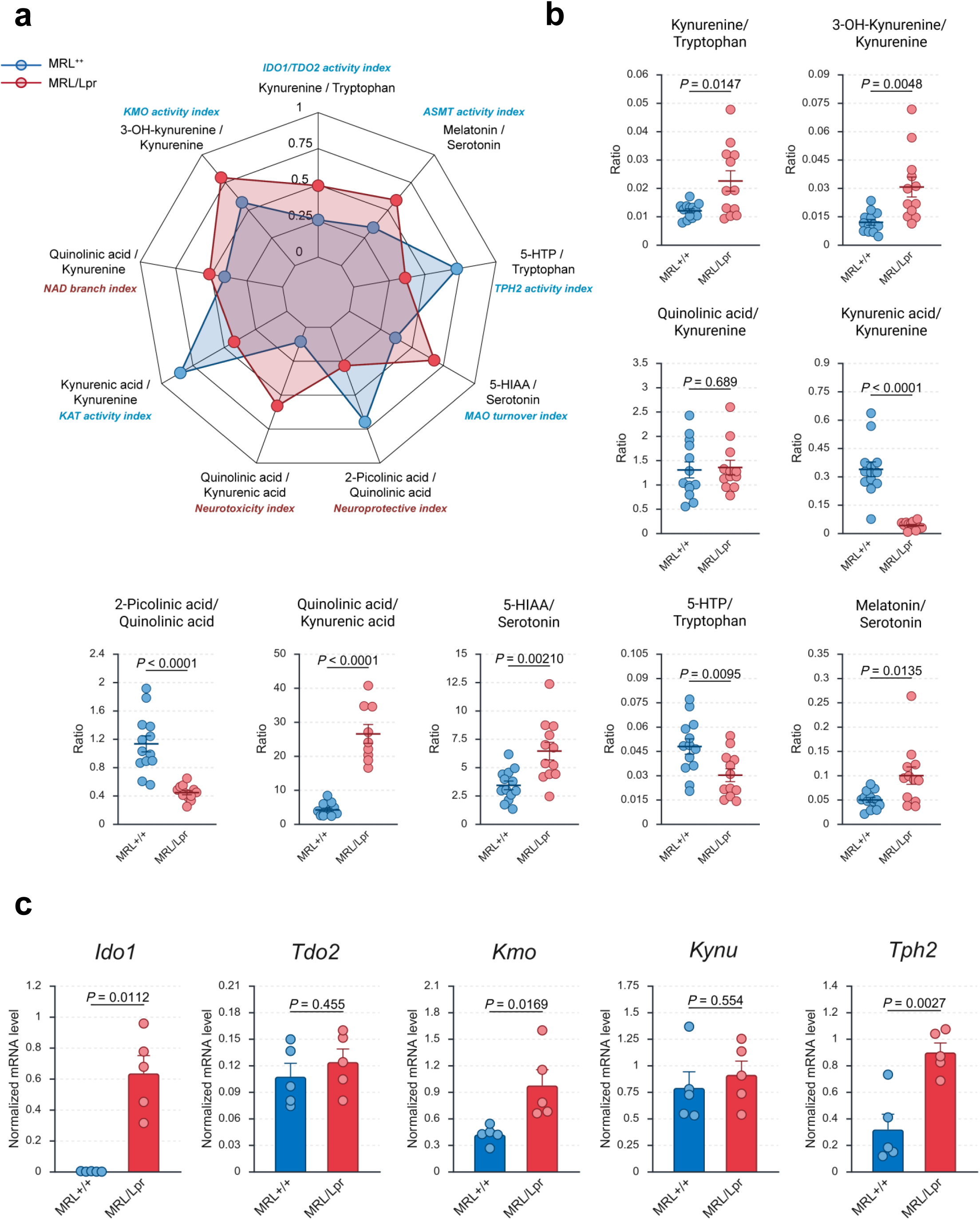
Metabolite ratios and enzyme expression reveal IFNγ-driven IDO1/KMO activation and impaired serotonin biosynthesis. **(A)** Radar plot of metabolite ratios used as functional activity indices: KYN/TRP (IDO1/TDO2 activity index), 3-HK/KYN (KMO activity index), Melatonin/Serotonin (ASMT activity index), 5-HTP/TRP (TPH2 activity index), 5-HIAA/Serotonin (MAO turnover index), QA/KA (neurotoxic index), PA/QA (neuroprotective index), QA/KYN (NAD branch index), and KA/KYN (KAT activity index). **(B)** Individual metabolite ratios in the frontal cortex, including KYN/TRP, 3-HK/KYN, 5-HTP/TRP, 5-HIAA/Serotonin, QA/KA, PA/QA, QA/KYN, and KA/KYN. **(C)** Normalized cortical mRNA expression levels (qPCR) of *Ido1*, *Tdo2*, *Kmo*, *Kynu*, and *Tph2*. Each point represents an individual mouse, reflecting the number of biological replicates per group. Bars indicate group mean ± SEM. Statistical analyses were performed using appropriate parametric or non-parametric tests depending on data distribution (see Methods for details). A *P* value < 0.05 was considered statistically significant. Abbreviations: *TRP, Tryptophan; 5-HTP, 5-Hydroxytryptophan; 5-HIAA, 5-Hydroxyindoleacetic acid; KYN, Kynurenine; 3-HK, 3-Hydroxykynurenine; KA, Kynurenic acid; QA, Quinolinic acid; PA, 2-Picolinic acid*.

We next examined the expression of genes encoding key enzymes in these pathways (Fig. 5C). *Ido1* was strongly upregulated in MRL/Lpr mice, consistent with IFNγ-driven induction of the KP, while *Tdo2* and *Kynu* remained unchanged. *Kmo* expression was significantly increased, reinforcing diversion of KYN into 3-HK and QA. In parallel, the serotonergic pathway displayed paradoxical signs of compensation: *Tph2*, the rate-limiting enzyme for serotonin biosynthesis, was significantly upregulated despite reduced serotonin levels, suggesting a feedback response to neurotransmitter depletion.

We then asked whether systemic IFNγ levels could be directly linked to these cortical changes. At the metabolic level (Fig. 6A), plasma IFNγ concentrations showed a trend toward a positive correlation with the KYN/TRP ratio (ρ = 0.40, *P* = 0.06) and, more strikingly, with the QA/KA ratio (ρ = 0.77, *P* < 0.001), demonstrating that systemic inflammation is tightly associated with a cortical metabolic profile favoring excitotoxic outputs. At the transcriptional level (Fig. 6B), IFNγ showed strong positive associations with *Ido1* (ρ = 0.82, *q* = 0.01), *Kmo* (ρ = 0.71, *q* = 0.046), and *Tph2* (ρ = 0.87, *q* = 0.01), reinforcing its role as a key upstream regulator of both KYN and serotonin metabolism.

**Figure 6:**
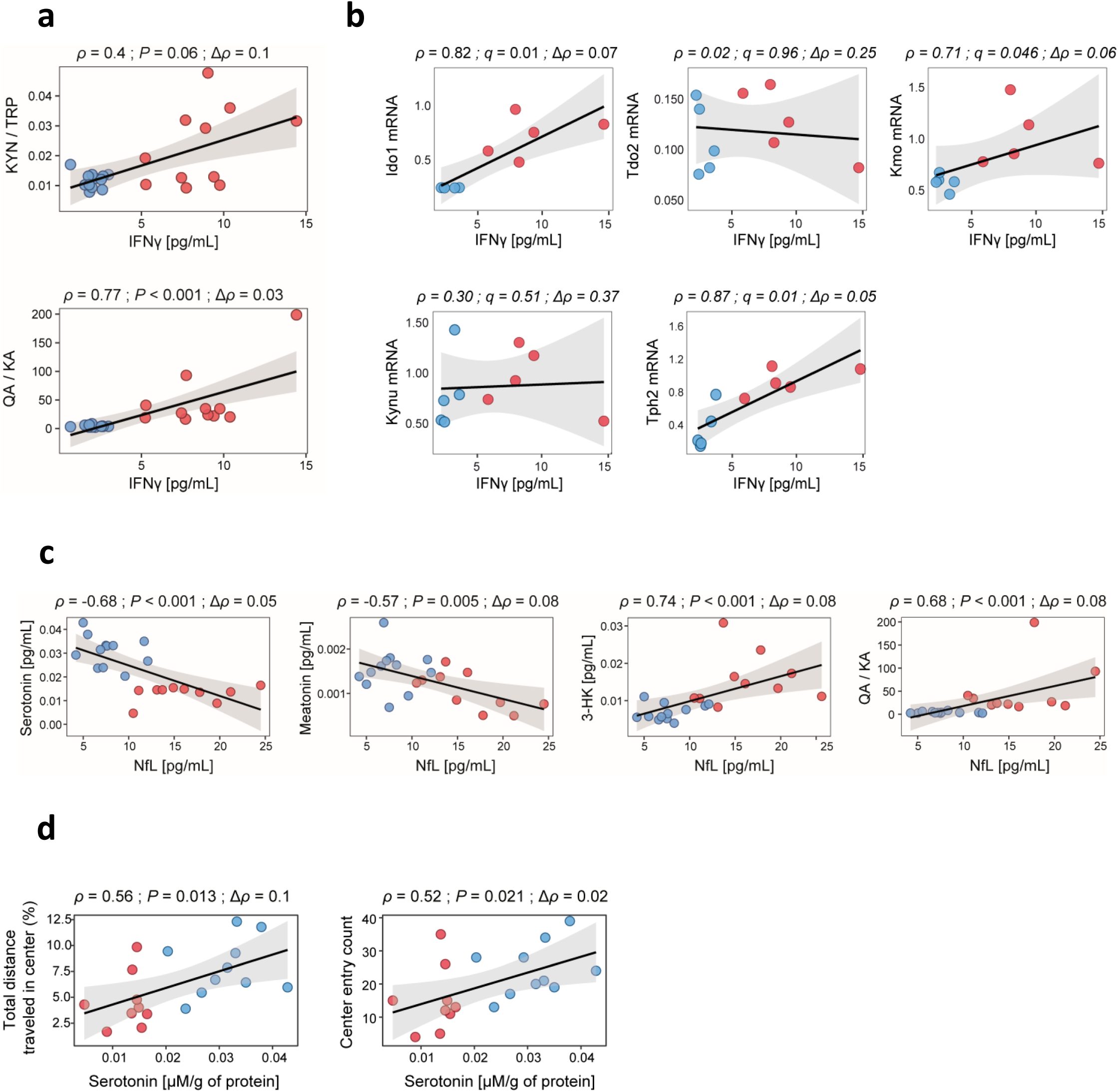
Systemic IFNγ connects tryptophan remodeling to NfL elevation and serotonergic-dependent anxiety-like behavior. (A–B) Correlations between plasma IFNγ concentrations (pg/mL) and frontal cortex metabolite ratios (KYN/TRP, QA/KA) or frontal cortex gene expression of key tryptophan metabolism enzymes measured by qPCR (*Ido1*, *Tdo2*, *Kmo*, *Kynu*, *Tph2*). **(C)** Correlations between plasma NfL concentrations (pg/mL) and cortical metabolites, including Serotonin, Melatonin, 3-HK, and the QA/KA ratio. **(D)** Correlations between cortical serotonin levels and behavioral performance in the open field test, including percentage of total distance traveled in the center and number of center entries. Each point represents an individual mouse, reflecting the number of biological replicates per group. Regression lines are shown with shaded 95% confidence intervals. Reported statistics include correlation coefficient (ρ), Δρ. *P* value or FDR-adjusted *q* value < 0.05 considered statistically significant. Abbreviations: *KYN, Kynurenine; TRP, Tryptophan; QA, Quinolinic acid; KA, Kynurenic acid; 3-HK, 3-Hydroxykynurenine; NfL, neurofilament light chain*.

Importantly, NfL, a sensitive biomarker of axonal injury, displayed highly significant associations with these pathways (Fig. 6C). Elevated plasma NfL levels correlated negatively with both serotonin and melatonin, and positively with 3-hydroxykynurenine (3-HK) and the quinolinic-to-kynurenic acid (QA/KA) ratio, reinforcing its relevance as a translational marker of metabolic imbalance and neuronal vulnerability. These associations indicate that NfL not only reflects ongoing neurodegeneration but also captures the metabolic shift toward excitotoxic KYN products.

Finally, we explored whether serotonergic deficits in the cortex were linked to the behavioral alterations observed in MRL/Lpr mice. As shown in Fig. 6D, cortical serotonin levels positively correlated with both the percentage of distance traveled in the center and the number of center entries in the open field, two measures commonly associated with reduced anxiety-like behavior. Thus, lower serotonin availability was directly associated with increased anxiety-like behavior, providing a functional bridge between systemic IFNγ, cortical metabolic reprogramming, axonal injury, and behavioral vulnerability.

Together, these findings demonstrate that systemic IFNγ acts as a central upstream driver that diverts TRP metabolism away from serotonin toward excitotoxic KYN products, in the Mrl/Lpr mouse model. This metabolic shift correlates with neuroaxonal injury (NfL) and translates into measurable behavioral deficits, thereby outlining an associative continuum from peripheral inflammation to metabolic imbalance, axonal injury, and behavioral dysfunction.

## Discussion

NPSLE remains a major clinical challenge, with its heterogeneous manifestations reflecting the interplay of systemic autoimmunity, CNS vulnerability, and complex immunometabolic mechanisms (36). Although inflammation is a defining hallmark of NPSLE, progress in characterizing the molecular links between systemic immune activation and CNS dysfunction remains insufficiently defined (37–39). Clinical studies consistently report elevated pro-inflammatory circulating cytokines (*e.g.*; IFNγ, TNFα, IL-6), together with neuroimaging evidence of hippocampal and cortical atrophy, and broad metabolic disturbances in plasma and CSF (30,40–43). Using the MRL/Lpr mouse model, a well-established experimental system for NPSLE, our integrative approach combined behavioral phenotyping, cytokine profiling, plasma NfL measurements, region-specific CNS cytokine expression, and targeted cortical metabolomics to delineate a mechanistic cascade linking systemic immune activation to neuronal injury and behavioral dysfunction. Several of these alterations, including behavioral deficits and TRP/KYN pathway imbalance, have previously been reported in the MRL/lpr model (27). While previous studies were mainly descriptive, here we extend these observations by systematically integrating NfL, a translational biomarker of axonal injury, together with immuno-metabolic readouts. This approach provides a more comprehensive view of the continuum from inflammation to neuronal vulnerability. It also extends prior observations by showing that IFNγ emerges as a key immune driver capable of reshaping cortical vulnerability through transcriptional and metabolic reprogramming of tryptophan catabolism, thereby establishing a mechanistic continuum from peripheral inflammation to metabolic imbalance, excitotoxicity, axonal injury, and behavioral vulnerability.

A first key observation is that MRL/Lpr mice exhibited the expected systemic features of lupus, including proteinuria, nephromegaly, and splenomegaly, but also clear neurobehavioral alterations. Anxiety-like behavior in the open field and memory impairments in cognitive tasks (Y-maze and novel object recognition) were prominent, highlighting functional consequences of systemic autoimmunity on CNS function. These alterations were paralleled by strong elevations of plasma NfL, a sensitive marker of axonal injury, which correlated with a coordinated pro-inflammatory cytokine program dominated by IFNγ, TNFα, IL-6, IL-1β, and IL-10. By contrast, IL-17A and GM-CSF remained unchanged, suggesting that, unlike MS or Th17-driven encephalitides, NPSLE is primarily associated with a Th1-oriented cytokine axis (44–46). PCA and composite scoring confirmed that these cytokines form an integrated systemic inflammatory load closely tied to NfL levels. These findings extend observations from MS, where NfL reflects inflammatory activity, and AD, where it tracks neurodegeneration, to NPSLE, supporting its potential use in combination with immune-derived indices as a biomarker framework for CNS involvement in lupus. While NfL has reached broad validation in MS and AD, its application in SLE remains to be firmly established, with current evidence promising but still limited. Together, our results strengthen the concept that immune dysregulation in lupus translates into neuronal vulnerability, and they position NfL as a promising translational biomarker for CNS involvement in SLE. While no histological validation of axonal injury was performed in this study, the strong and consistent correlations with cytokines and metabolic indices support its biological relevance in this context (47–50).

Region-specific analyses of CNS cytokine expression further revealed a striking mosaic of local immune signatures. IL-1β dominated in the hippocampus, IL-6 in the cerebellum, IFNγ in the frontal cortex, and both IFNγ and IL-10 in the spinal cord. Interestingly, IL-10, a key anti-inflammatory cytokine, was reduced in the hippocampus despite being elevated systemically, pointing to a selective loss of protective regulation in a region already known to be vulnerable in NPSLE. These findings echo neuroimaging evidence highlighting hippocampal and cortical involvement in patients, and emphasize the importance of considering local immune environments when interpreting CNS pathology (51–54). Such region-specific cytokine patterns have been described in MS and viral encephalitis, where focal immune microenvironments underlie selective vulnerability (55,56). The strong pro-inflammatory signatures detected in the cerebellum and spinal cord, together with increased spinal cord mass, raise the possibility that focal neuroinflammation may drive hypertrophic or gliotic changes in lupus, analogous to transient spinal cord enlargement reported in MS before atrophy (57–59). This expands the view that lupus-related neuroinflammation may extend well beyond hippocampal and cortical circuits, and underscores the value of regionally resolved analyses for disentangling clinical heterogeneity.

The dominance of IFNγ in the frontal cortex is particularly compelling, as it provides a mechanistic link to downstream metabolic remodeling of the tryptophan pathway. Cortical metabolomics revealed preserved precursor levels but a profound redirection of flux away from serotonin and melatonin biosynthesis toward the KP. This shift was marked by reductions in serotonin, melatonin, 5-HTP, 5-HIAA, KA, and PA, alongside increases in KYN, 3-HK, and QA, coupled with transcriptional upregulation of *Ido1* and *Kmo*. Ratios such as KYN/TRP and QA/KA capture this imbalance, with the latter serving as a widely recognized marker of excitotoxic potential (Weissman-Tsukamoto et al. 2025). The pronounced rise in QA/KA mirrors clinical observations of elevated CSF QA/KA in NPSLE patients, where it correlates with cognitive dysfunction, depression, and fatigue (60,61). These results are also consistent with prior work in the MRL/Lpr model that documented depression-like behavior, visuospatial memory deficits, and increased cortical and hippocampal QA, together with elevated QA/KA ratios (27). Our findings confirm these neurochemical alterations and extend them by linking cortical metabolic remodeling to systemic IFNγ, neuronal injury (NfL), and behavioral alterations.

Indeed, plasma IFNγ strongly correlated with QA/KA ratios and the expression of *Ido1*, *Kmo*, and *Tph2*, consolidating its role as a central upstream regulator of both KYN and serotonin metabolism. The observed *Tph2* upregulation may reflect a compensatory attempt to restore serotonin synthesis in the face of neurotransmitter depletion. Importantly, NfL levels were inversely associated with serotonin and melatonin, and positively with 3-HK and QA/KA, supporting its utility as a translational biomarker of excitotoxic metabolic imbalance.

Finally, cortical serotonin levels correlated with exploratory behavior, aligning serotonergic depletion with anxiety-like phenotypes. While unlikely to be the sole driver, serotonergic imbalance emerges as a key mechanistic contributor to neuropsychiatric vulnerability in this model. Serotonin plays a central role in mood and anxiety regulation, particularly within prefrontal–limbic circuits, and its depletion has long been associated with heightened anxiety and depressive behaviors across species (62,63). Clinical studies similarly report reduced serotonergic metabolites in plasma and CSF associated with depression and anxiety in autoimmune and neuropsychiatric conditions (64,65). The convergence of IFNγ-driven tryptophan diversion, cortical serotonin depletion, and measurable behavioral alterations in MRL/Lpr mice therefore supports the broader concept that immune–metabolic dysregulation can directly shape affective behavior.

While our findings provide novel mechanistic insights, several limitations warrant acknowledgement. Analyses were restricted to female mice at a single disease stage (17 weeks), chosen to maximize detection of inflammatory and metabolic alterations. Future studies including males and longitudinal designs will be necessary to assess temporal and sex-related dynamics. In addition, although correlations were strong and robust to leave-one-out testing, they remain associative, and interventional studies directly targeting IFNγ, IDO1, or KMO will be required to establish causality. Moreover, the MRL/Lpr strain does not fully recapitulate all pathogenic mechanisms of human SLE, as disease progression in this model is primarily IFNγ–driven rather than IFNα–driven and strongly influenced by the Fas receptor mutation, which has no direct counterpart in most human cases (12). Nevertheless, since IFNα is also a potent inducer of tryptophan catabolism through IDO1 and KMO, the metabolic pathways identified here likely remain relevant to human lupus. Finally, metabolomic analyses were limited to the frontal cortex, selected for its central role in cognition and affective regulation (66–68). Extending these analyses to hippocampus, cerebellum, and spinal cord, regions with distinct inflammatory signatures, will be crucial for fully mapping region-specific vulnerability.

In summary, our study identifies systemic IFNγ as a central upstream driver that links peripheral autoimmunity to cortical metabolic remodeling and axonal injury in lupus-prone mice. We propose an integrative cascade in which immune activation is associated with serotonergic depletion, KYN excitotoxicity, NfL elevation, and behavioral alterations. More broadly, these findings highlight the emerging immune–metabolic crosstalk as a key determinant of brain vulnerability across neuropsychiatric and neurodegenerative disorders, pointing to opportunities for biomarker development and mechanism-based therapeutic interventions, while underscoring the need for validation in clinical cohorts.

## List of abbreviations

3-HK: 3-Hydroxykynurenine
5-HIAA: 5-Hydroxyindoleacetic acid
5-HTP: 5-Hydroxytryptophan
AD: Alzheimer’s disease
CNS: Central nervous system
CSF: Cerebrospinal fluid FDR False discovery rate
GM-CSF: Granulocyte–macrophage colony-stimulating factor
IDO1: Indoleamine 2,3-dioxygenase-1
IFNγ: Interferon-gamma
IL-1β: Interleukin-1 beta
IL-6: Interleukin-6
IL-10: nterleukin-10
IL-17A: Interleukin-17A
KA: Kynurenic acid
KAT: Kynurenine aminotransferase
KMO: Kynurenine 3-monooxygenase
KP: Kynurenine pathway
KYN: Kynurenine
Kynu: Kynureninase
MAO: Monoamine oxidase
MS: Multiple sclerosis
NfL: Neurofilament light chain
NOR: Novel Object Recognition
NPSLE: Neuropsychiatric systemic lupus erythematosus
OF: Open Field
PA: 2-Picolinic acid
PCA: Principal component analysis
QA: Quinolinic acid
RI: Recognition index
ROC: Receiver operating characteristic
SAB: Spontaneous alternation behavior
SLE: Systemic lupus erythematosus
Tdo2: Tryptophan 2,3-dioxygenase
Th1: Type 1 helper T cell
TNFα: Tumor necrosis factor alpha
Tph2: Tryptophan hydroxylase 2
TRP: Tryptophan

## Acknowledgements

We thank Christian Klein for his valuable technical assistance.

## Authors’ contributions

KM: Conceptualization; Formal analysis; Investigation; Data curation; Visualization; Writing – original draft; Writing – review & editing. RMGR: Formal analysis; Writing – review & editing. ED: Resources (behavioral testing); Writing – review & editing. OB: Formal analysis; Writing – review & editing. JMA: Formal analysis; Writing – review & editing. JLG: Formal analysis; Writing – review & editing. AGMN: Funding acquisition; Writing – review & editing. HJD: Conceptualization; Investigation; Writing – original draft; Writing – review & editing.

## Data availability statement

All data supporting the findings of this study are available from the corresponding author upon reasonable request.

## Supplementary Material

Supplementary Material S1

